# Physical confinement promotes mesenchymal trans-differentiation of invading transformed cells *in vivo*

**DOI:** 10.1101/2022.07.21.501012

**Authors:** Teresa Zulueta-Coarasa, John Fadul, Marjana Ahmed, Jody Rosenblatt

## Abstract

Metastasis is tightly linked with poor cancer prognosis, yet it is not clear how transformed cells become invasive carcinomas. We previously discovered that single KRas^V12^-transformed cells can invade directly from the epithelium by basal cell extrusion. During this process, cells de-differentiate by mechanically pinching off their epithelial determinants, but how they trans-differentiate into a migratory, mesenchymal phenotype is not known. Here, we demonstrate that basally extruded KRas^V12^-expressing cells become significantly deformed as they invade the zebrafish body. Decreasing the confinement that cells experience after they invade reduces the percentage of KRas^V12^ cells that trans-differentiate into mesenchymal cell types, while higher confinement increases this percentage. Additionally, increased confinement promotes accumulation of internal masses over time. Altogether, our results suggest that mechanical forces drive not only de-differentiation of KRas^V12^-transformed epithelial cells as they invade but also contribute to their re-differentiation into mesenchymal phenotypes that contribute to distant metastases.

## INTRODUCTION

Although metastasis is the predominant cause of mortality in cancer patients, how tumour cells invade to form metastases is not well understood. For a cancer cell to metastasize, it must first invade from epithelia, where most solid tumours originate, and then trans-differentiate to acquire a malignant phenotype. The prevailing metastasis model suggests that as cells accumulate mutations, they first form primary masses, which they later escape by downregulating epithelial specific genes to undergo an <u>E</u>pithelial to <u>M</u>esenchymal <u>T</u>ransition (EMT) [1]. Transition from epithelia to mesenchymal phenotypes allow cells to dissociate from the primary tumour so that they can invade and colonise distant organs. Because these invading cells have also stem-like qualities and markers that allow them to proliferate, survive, and transform into a spectrum of different cell types, the term <u>E</u>pithelial to <u>M</u>esenchymal Plasticity (EMP) has become a better descriptor and more widely used [2].

We have shown that oncogenic mutations in KRas that drive aggressive tumours [3-5] induce invasion of transformed epithelial cells by hijacking cell extrusion [6,7], a process that epithelia normally use to promote cell death [8]. In epithelial cell extrusion, a basal intercellular actomyosin cable contracts to squeeze one cell apically from the layer to die [8,9]. KRas mutations disrupt apical extrusion, causing cells to accumulate into masses or aberrantly extrude basally, underneath the epithelium [6,7]. Importantly, using zebrafish epidermis as a model for simple epithelia where carcinomas form, we previously found that transformed cells basally extrude directly from the epithelium, at sites separate from where they form masses [7]. <u>B</u>asal <u>c</u>ell <u>e</u>xtrusion (BCE) enables KRas-transformed cells to invade, divide and migrate throughout the zebrafish body. While most invading cells die, if they lack a functional copy of the tumour suppressor p53, a commonly collaborating mutation in aggressive cancers, they can instead survive to form large internal masses [7]. Importantly for this study, some invading KRas^V12^-expressing cells trans-differentiate into mesenchymal phenotypes, with a smaller, but significant fraction adopting a neuronal-like morphology [7]. The mechanisms that regulate differential fates of invaded cells are unknown.

In contrast to previous EMT models, we previously found that de-differentiation of invading transformed epithelial cells occurs suddenly and mechanically as it invades by BCE by pinching off the apical membrane containing E-cadherin and other proteins essential for epithelial identity and function [7]. Whereas all basally extruded cells lack E-cadherin, only a fraction express the mesenchymal marker N-cadherin, suggesting a two-step model for EMT whereby cells dedifferentiate by BCE and some trans-differentiate into mesenchymal cell types via another mechanism [7]. Given that cells can invade directly from the epithelium, suggesting that the primary tumour is unlikely to influence the mesenchymal trans-differentiation. However, mechanical stress can induce EMT during embryonic development and tumorigenesis [10,11], suggesting a potential role for mechanical strain promoting mesenchymal transformed cell differentiation as they migrate throughout the tight confines of extracellular matrix and organs within the body. Here, we experimentally alter the microenvironment that cells expressing EG-FP-KRas^V12^ encounter as they invade and migrate throughout the body of zebrafish to test the role of mechanical strain on their fates.

## RESULTS

### Transformed cells deform following invasion

To investigate the role of the mechanical microenvironment in promoting cell invasion we injected zebrafish embryos with DNA to mosaically express krt4:EGFP-KRas^V12^ in the outer epidermal layer along with a p53 morpholino (MO) to enable invading cell survival. Unless otherwise indicated, all our experiments were quantified at 2 days post-fertilisation (dpf). To determine if the tight confinement that invading cells encounter causes mechanical deformation of EGFP-KRas^V12^ cells, we measured the circularity of their nuclei before and after BCE in embryos expressing mCherry fused to the actin-binding domain of Utrophin [12] along with H2B-RFP mRNA to visualize nuclei [13]. We found that the circularity of the nucleus in KRas^V12^-expressing cells decreases sharply after BCE (Fig. 1a-c and Supplemental Movie 1). Furthermore, the nuclear circularity of invaded KRas^V12^ cells was significantly lower than in those remaining within the embryonic surface, irrespective of their location within the body. (0.083±0.01 versus 0.094±0.01, *P* < 0.001, *P* < 0.01, Fig. 1d-g). The changes in nuclear deformation following invasion suggests that invading cells become confined as they migrate through the dense tissue of the zebrafish body. When cancer cells migrate through confined spaces, nuclear deformation can result in nuclear envelope rupture and DNA damage [14-17]. Moreover, compression-derived DNA damage in breast cancer cells results in a snail1-dependent invasive phenotype [18]. We sought to investigate if nuclear deformation results in DNA damage in invading EGFP-KRas^V12^ cells by immunostaining with the marker phospho-Histone H2A.X. We found that only 1.3 ± 1.3% of KRas^V12^-transformed cells in the surface of the embryo presented nuclear damage, while 12.23 ± 2.4% of invaded KRas^V12^-positive cells do so (*P* < 0.05, Fig. 2h-j). These data suggest that nuclear deformation cells experience during invasion and migration can damage their nuclei.

**Fig. 1:**
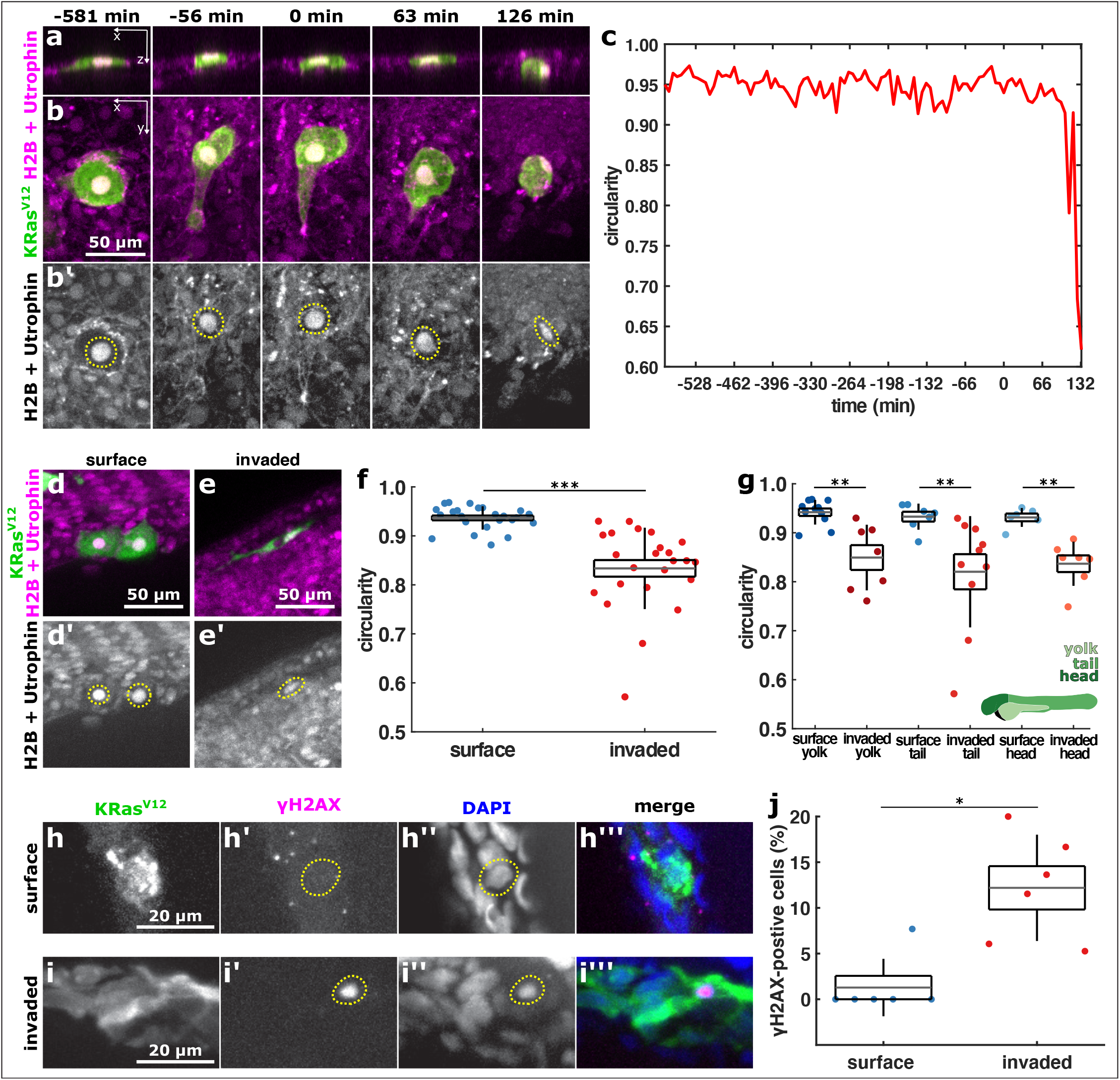
KRas^V12^ cell nuclei deform after they invade. a, b, orthogonal and projection views of an invading cell expressing EGFP-KRas^V12^ (green in a, b) and H2B-RFP and mCherry-Utrophin (b’, both, magenta in a, b) in XZ (a) or XY (b). Time before and after BCE is indicated. c, nuclear circularity over time for the cell shown in a, b. d, e, example cells expressing EGFP-KRas^V12^ (green in d, e) and H2B-RFP and mCherry-Utrophin (d’, e’ magenta in d, e) in the outer epithelium (d) or invaded inside the embryo (e), quantified in f & g. f, nuclear circularity in surface (n = 25) and invaded (n = 24) KRas^V12^ cells. g, nuclear circularity in surface and invaded KRas^V12^ cells located in the yolk (n _surface_ = 11 and n_invaded_ = 7), the tail (n_surface_ = 7 and n_invaded_ = 10) and the head (n_surface_ = 7 and n_invaded_ = 7) of the embryo. The embryo diagram in green denotes how the different regions were classified. h, EGFP-KRas^V12^ cells in an embryo stained with GFP (h, h’’’), phospho-histone H2A.X (h’, h’’’) and DAPI (h’’, h’’’). In a, b, d, e, the yellow dotted lines outline the cell nucleus. In f, g, error bars are s.d., boxes are s.e.m., and grey lines denote the mean. **P < 0.01, ***P < 0.001.

**Fig. 2:**
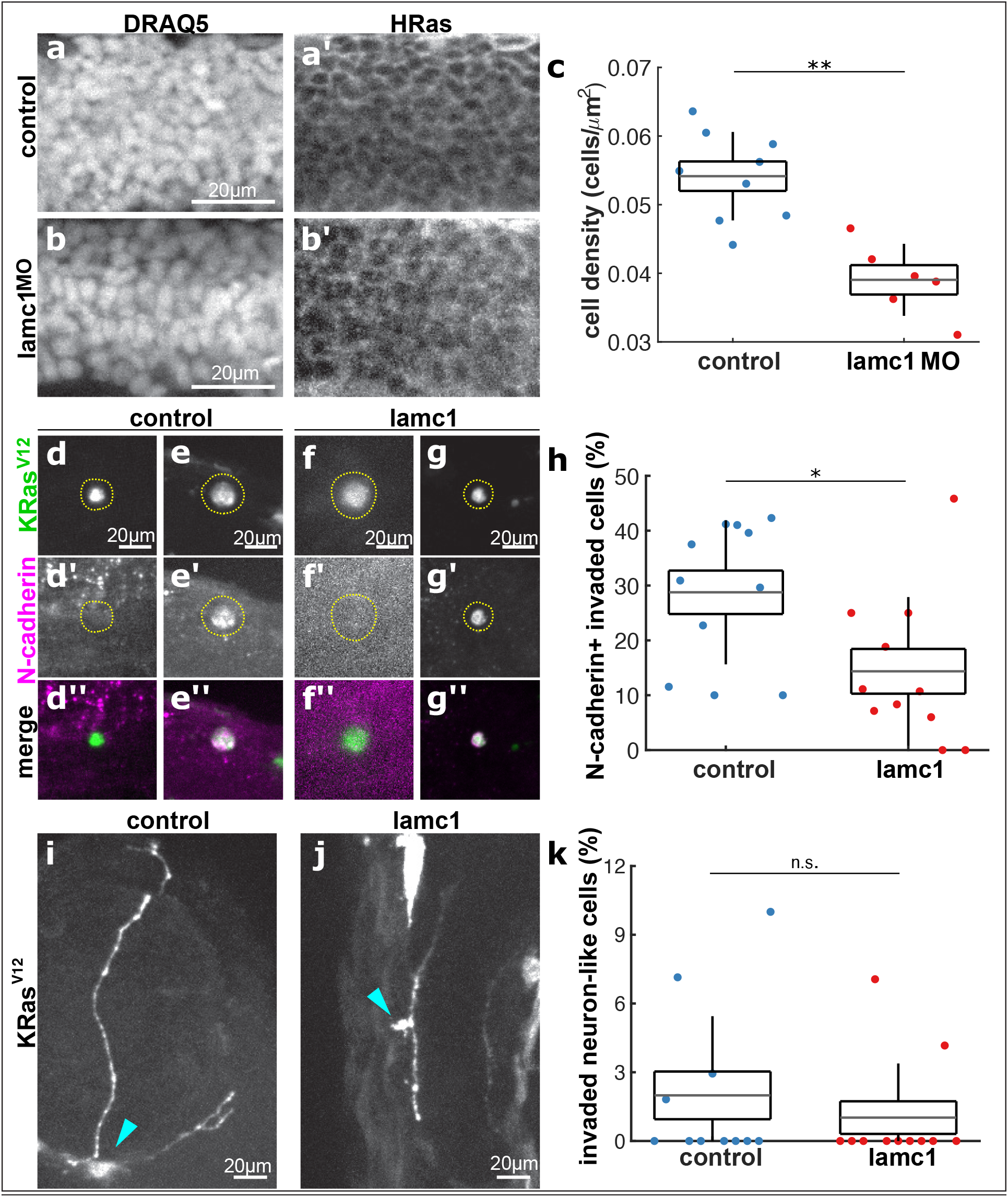
Loss of laminin reduces KRasV12 cell trans-differentiation into mesenchymal phenotypes. a, b, neural tube cells expressing HRas-EGFP in control (a) and lamc1 MO-injected (b) embryos, stained with DRAQ5 (a, b) and GFP (a’, b’) used for cell density quantification in c, where control embryos are blue (n = 9) and lamc1 morphants are red (n = 6). d-g, images of EGFP-KRasV12-expressing invaded cells in controls (d, e) and lamc1 mutants (f, g) stained with GFP (d, e, f, g) and N-cadherin (d’, e’, f’, g’), where the yellow dotted lines outline cells of interest, quantified in h, as percentage of KRasV12 invaded cells expressing N-cadherin in control (n = 11) and lamc1 mutants (n = 11). i, j, neuron-like morphology cells expressing EG-FP-KRasV12 in control (i) and lamc1 mutants (j), where cyan arrowheads denote cell bodies, quantified in k, as percentage of KRasV12 invaded cells with neuron-like morphologies in control (n = 11) and lamc1 mutants (n = 11). In c, h, k, the error bars are s.d., the box s.e.m., and the grey lines denote the mean. n.s., not significant, *P < 0.05, **P < 0.01.

### Reduced ECM density results in fewer KRas^V12^ mesenchymal cells

Since we previously found that invading cells could differentiate into mesenchymal and neuronal-like cells following invasion [7], we next tested if tissue confinement could impact either cell fate. To do so, we tested whether altering the physical environment that EGFP-KRas^V12^ invading cells encounter affects their trans-differentiation into different cell types. To reduce the extracellular matrix density within embryos [19], we disrupted a key ECM component, laminin, using *lamc1* mutants that produce no detectable laminin [20] or *lamc1* morpholinos that significantly reduce laminin expression [21]. *Lamc1* mutants have more apoptotic cells in the trunk [21], which could further reduce cell density and confinement. To assess if laminin loss results in reduced overall cell density, we quantified neural tube cell density of control versus *lamc1* MO embryos using HRas-EGFP to define cell outlines and DRAQ5 to visualize cell nuclei (Fig. 2a, b). We found that *lamc1* morphants had 28% reduced neural tube cell density compared to controls (*P* < 0.01, Fig. 2c). To investigate if decreasing cell density and confinement affects trans-differentiation, we stained control and *lamc1* mutant embryos with the mesenchymal marker N-cadherin. While 28.8 ± 4.0% of invaded KRas^V12^/ p53MO cells in control embryos express N-cadherin, only 14.4 ± 4.1% of cells invading in *lamc1* mutants do so (*P* < 0.05, Fig. 2d-h). Similarly, *lamc1* morphants reduced the number of N-cadherin-positive KRas^V12^ cells by 77%, compared to controls (*P* < 0.05, Fig. S1a-e). Interestingly, reduced confinement did not significantly alter the proportion of invaded KRas^V12^/ p53MO cells that adopt a neuron-like morphologies: 1.1 ± 0.7 % in lamc1 mutants compared to 2.0 ± 1.1 % in controls (*P* > 0.05, Fig. 2i-k). These data suggest that an intact ECM contributes to trans-differentiation of transformed cells into a mesenchymal but not neuron-like phenotypes.

### Increased confinement promotes trans-differentiation of mesenchymal cell types

Because decreasing the ECM density could affect cell fate by altering signalling via reduced ECM-cell adhesion, we manipulated the mechanical microenvironment independently of ECM composition.

**Fig. S1:**
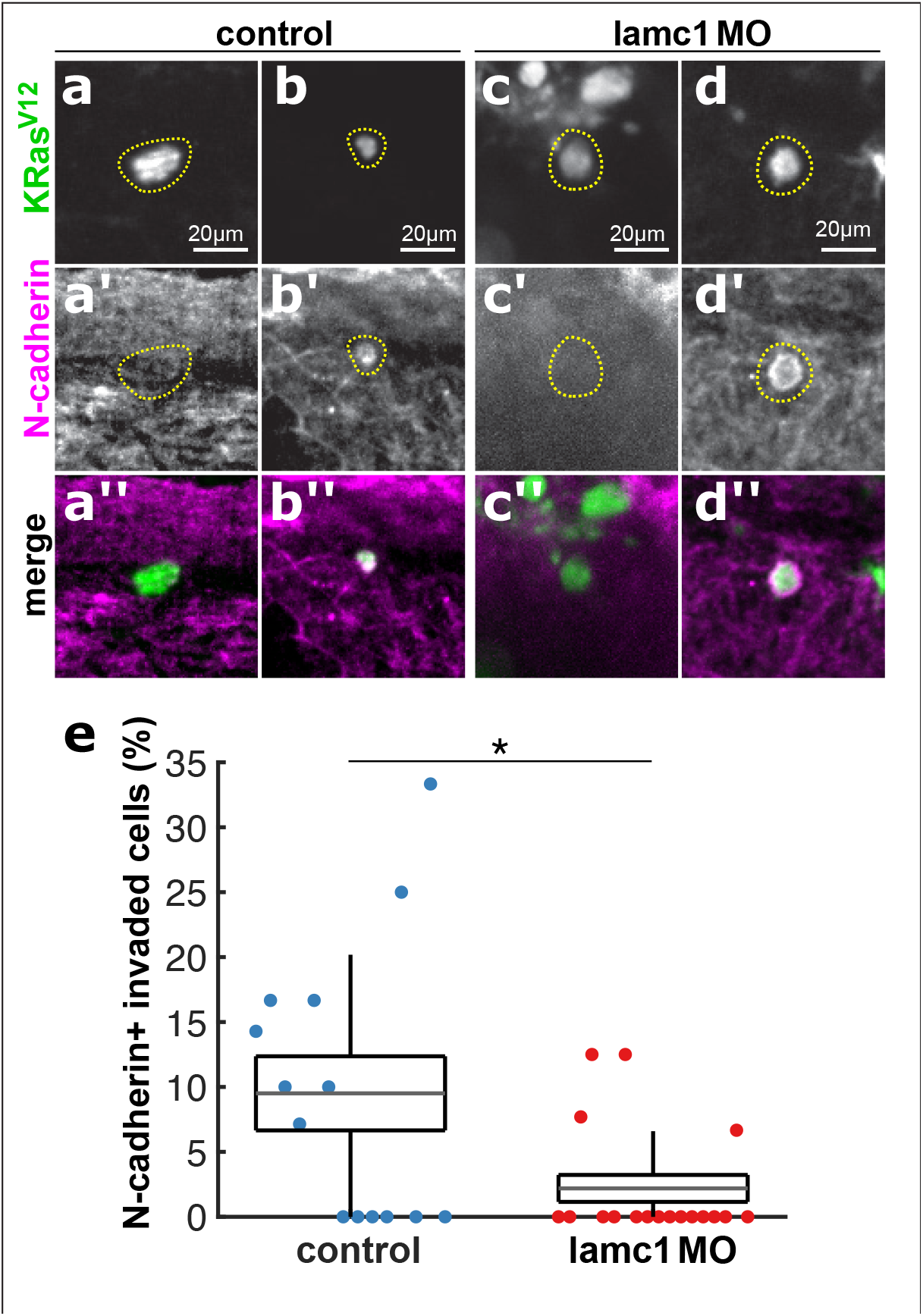
KRasV12 cells in lamc1 morphants present lower rates of trans-differentiation. a-d, EGFP-KRas^V12^-expressing invaded cells in controls (a, b) and lamc1 morphants (c, d) stained with GFP (a-d) and N-cadherin (a’-d’). The yellow dotted lines outline the cells of interest. e, percentage of KRas^V12^ invaded cells expressing N-cadherin in control (n = 20) and lamc1 morphants (n = 19).

To increase physical confinement experienced by KRas^V12^-expressing cells following invasion, we embedded embryos from 1 to 2 dpf in 2% agarose to compress them. Agarose-based confinement resulted in significantly smaller embryos, where the total embryo size (area) in 2% agarose (physical confinement) was 0.53 ± 0.01 mm^2^ (Fig. 3b, c) compared to 0.88 ± 0.03 mm^2^ in 0% agarose (unrestricted growth, *P* < 0.001, Fig. 3a, c). Embryos grown in 2% agarose also developed crooked tails (Fig. 3b) and thinner somites along the anterior-posterior axis (Fig 3d, e), suggesting that agarose compressed the tissue during embryonic development. To determine if smaller embryos result from compression or from defects in cell proliferation, we analysed the cell density of somites in embryos raised in 0 versus 2% agarose in zebrafish expressing HRas-EGFP for cell boundaries and stained with DRAQ5 to visualise cell nuclei. Development in 2% agarose increased the somite cell density by 22%, compared to control embryos (Fig. 3d-f, P < 0.05), suggesting that agarose causes physical compression, rather than reduced proliferation. To test if increasing confinement affects invading KRas-transformed cell fate, we compared their trans-differentiation into mesenchymal and neuronal-like cells in embryos developing in 0% agarose and in 2% agarose. 26.0 ± 2.4% of KRas^V12^/ p53MO invaded cells within embryos grown in 0% agarose expressed N-cadherin (Fig. 3g, h, k), whereas 38.8 ± 3.7 % did so in embryos grown in 2% agarose (*P* < 0.05, Fig. 3i-k). However, no significant differences in the percentage of KRas^V12^-positive neuronal-like cells were detected between embryos treated with 0% (6.3 ± 1.0%) and 2% agarose (3.9 ± 0.9%, *P* > 0.05, Fig. 3l-n). Together, these results suggest that mechanical compression promotes trans-differentiation of KRas^V12^-invaded cells into mesenchymal but not neuronal cell types.

**Fig. 3:**
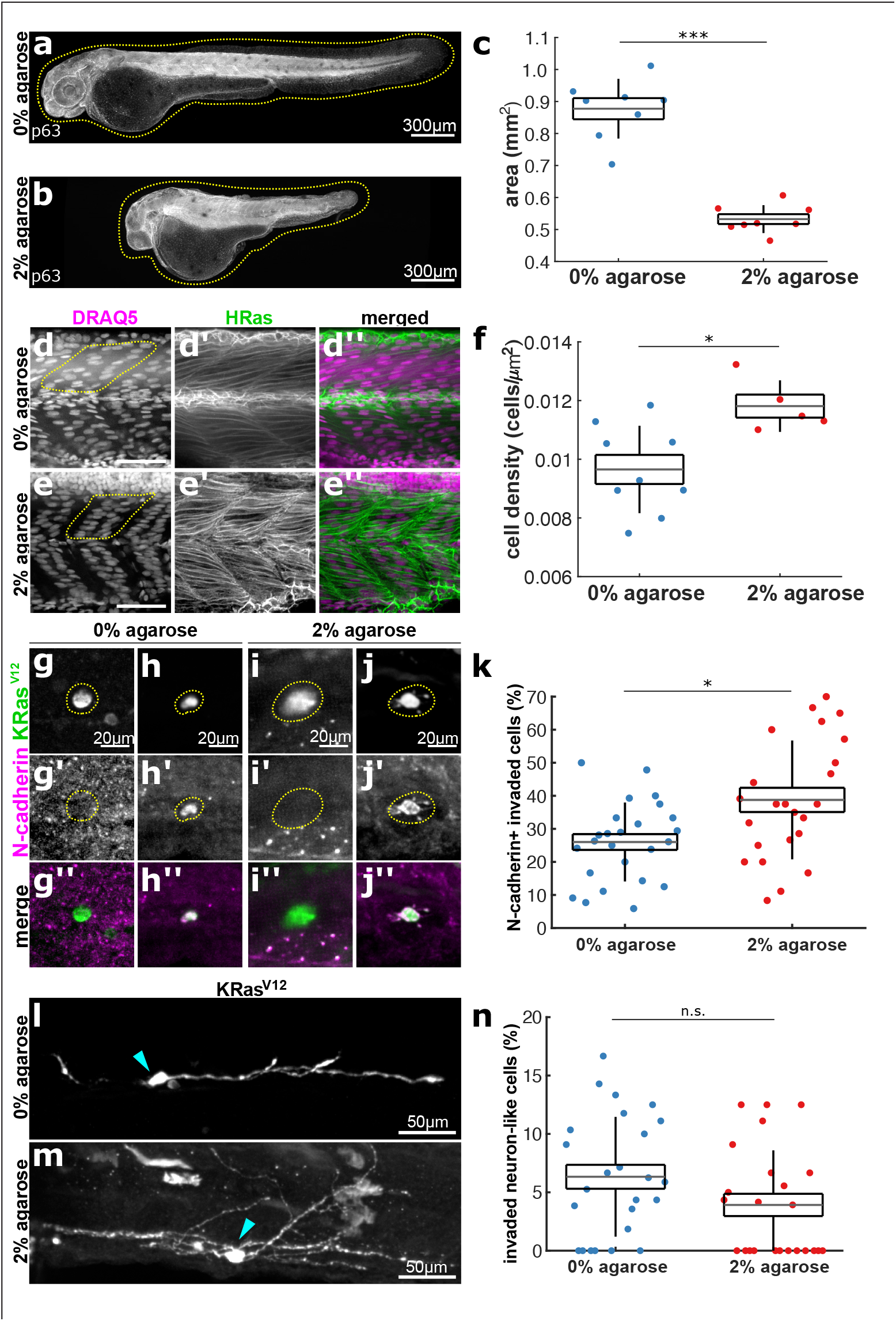
Embryo confinement in agarose promotes KRasV12 cell trans-differentiation into mesenchymal phenotypes. a, b, 2 dpf embryos grown from 1 to 2 dpf in 0% (a) or 2% agarose (b), with yellow dotted lines outlining embryos. c, area of 2 dpf embryos embedded in 0% (n = 8) or 2% agarose (n = 8) from 1 to 2 dpf. d, e, somite cells expressing HRas-EGFP in 0% (d) and 2% (e) agarose-treated embryos, stained with DRAQ5 (d, e, d’’, e’’) and GFP (d’, e’, d’’, e’’). The yellow dotted lines denote half somites. f, somite cell densities in control (0% agarose) embryos (blue, n = 9) and embryos grown in 2% agarose (red, n = 5). g-j, invaded EGFP-KRasV12-expressing mesenchymal cells in embryos embedded in 0% (g, h) and 2% (I j) agarose stained with GFP (g-j, g’’-j’’) and N-cadherin (g’-j’, g’’-j’’). Yellow dotted lines outline cells of interest. k, percentage of KRasV12 invaded cells expressing N-cadherin in 0% (n = 25) and 2% (n = 24) agarose embryos. l, m, cells expressing EG-FP-KRasV12 stained with GFP with a neuron-like shape in embryos grown in 0% (l) and 2% (m) agarose. Arrowheads denote the cell body. n, % of KRasV12 invaded cells adopting neuron-like morphologies in 0% (n = 25) and 2% (n = 24) agarose. In c, f, k, n, the error bars show the s.d., the box indicates the s.e.m. and the grey lines denote the mean. n.s., not significant, *P < 0.05, ***P < 0.001..

### Altering confinement affects internal mass formation

Embryos injected with KRas^V12^ in a *p53* mutant background develop internalized cell masses by 4.5 dpf, typically within the head [7]. The greater percentage of KRas^V12^-expressing mesenchymal cells caused by increased confinement, could result in more or bigger masses by 4.5 dpf. However, compression could also cause invading cells to die instead of contributing to these masses. To investigate if mechanical induction of trans-differentiation affects the formation of internal masses, we embedded KRas^V12^/ p53MO-injected embryos in 0% or 2% agarose from 1 to 2 dpf and fixed them at 4.5 dpf. We found that both types of embryos developed cell masses underneath the E-cadherin-stained epithelium (Fig. 4a, b). Importantly, embryos grown in high confinement developed more internal masses than embryos grown in 0% agarose (1.4 ± 0.3 vs. 0.6 ± 0.2 masses per embryo respectively, *P* < 0.05, Fig. 4a-c). However, the size of the masses remained unaffected in 2% agarose embryos (0.006 ± 6.08 × 10^−4^ mm^2^) compared to controls (0.006 ± 8.05 × 10^−4^ mm^2^, *P* > 0.05, Fig. 4d). These data suggest that mechanically increasing the percentage of KRas^V12^-expressing cells that trans-differentiate into mesenchymal phenotypes results in more internal masses by 4.5 dpf. To test if reducing confinement affects the later development of internal masses, we compared the number of cell masses in control KRas^V12^/ p53MO-injected embryos, and embryos also injected with *lamc1* MO. *Lamc1* morphants developed fewer internal cell masses than control embryos (0.2 ± 0.1 vs. 1.3 ± 0.3 masses per embryo respectively, *P* < 0.05, Fig. 4e-g). Although there were fewer masses within *lamc1* morphants, the sizes of these masses did not vary to those seen in controls (0.006 ± 3.2 × 10^−3^ mm^2^ in lamc1 to 0.005 ± 5.81 × 10^−4^ mm^2^ in controls *P* > 0.05, Fig. 4h). However, the low number of masses in *lamc1* morphants make statistical comparison difficult. Altogether, these results suggest that confinement has a role in the formation of internal cell masses after invasion.

**Fig. 4:**
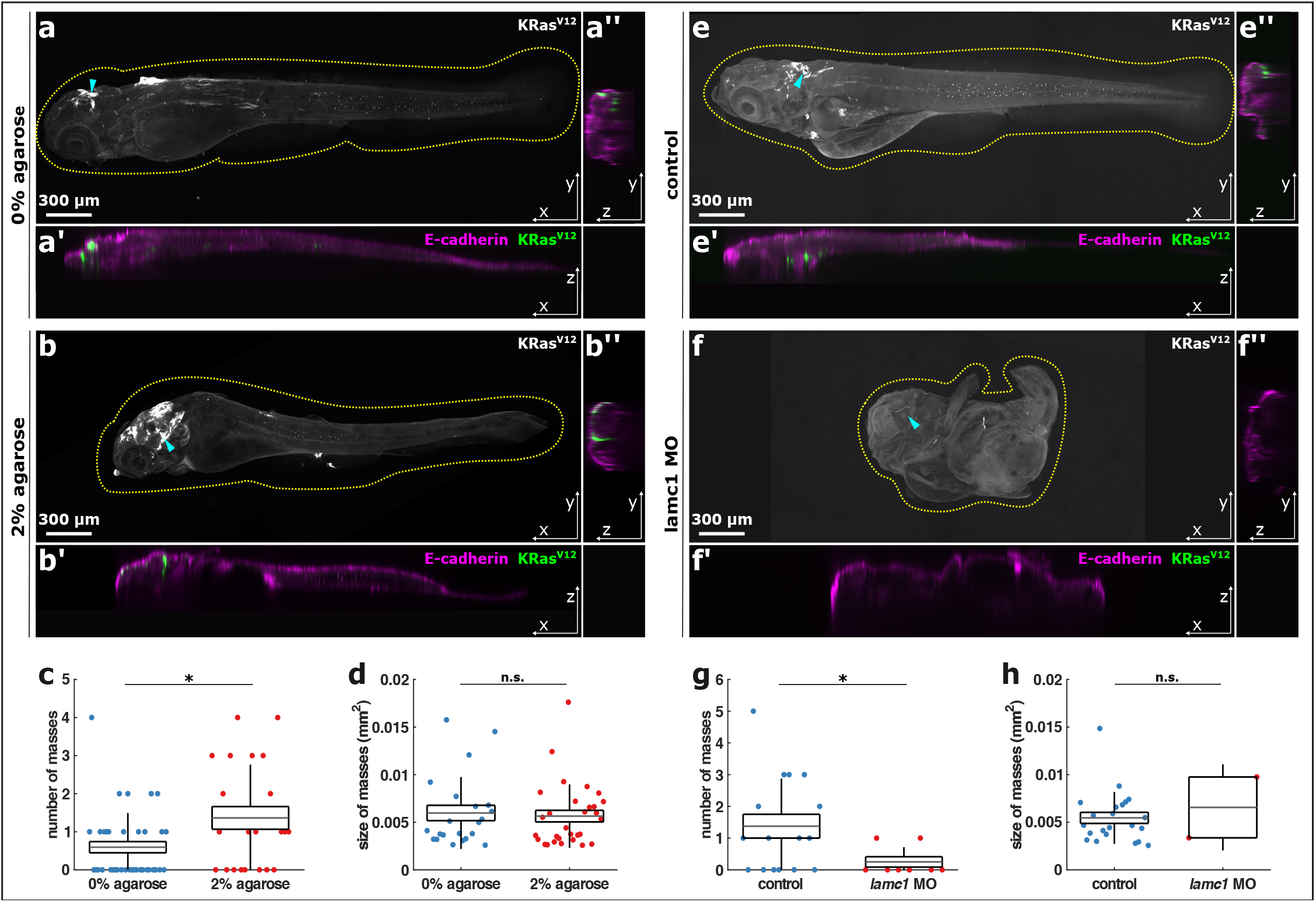
Experimental confinement promotes internal cell masses. a, b, representative orthogonal views of 4.5 dpf embryos quantified in c & d expressing EGFP-KRas^V12^, grown from 1 to 2 dpf in 0% (a) or 2% agarose (b), in XY (a, b), XZ (a’, b’) and YZ (a’’, b’’). Embryos were stained with E-cadherin to visualise the embryonic surface (magenta in a’-a’’ and b’-b’’) and GFP to visualise KRas^V12^ cells (a, b, green in a’-a’’ and b’-b’’). Yellow dotted lines outline the embryos and cyan arrowheads mark the coordinates in XY of orthogonal sections. Scale bars, 300 µm. c, number of cell masses per embryo in 0% (blue, n = 37) or 2% agarose (red, n = 22) embryos. d, cell mass areas from embryos grown in 0% (n = 22) or 2% agarose (n = 30). 4.5 dpf control (e) and lamc1 MO-injected (f) embryos expressing EGFP-KRas^V12^, as projections (e, f), XZ (e’, f’) and YZ sections (e’’, f’’). Yellow dotted lines outline embryos and arrowheads mark where orthogonal sections where taken. Scale bars, 300 µm. g, number of cell masses per embryo in controls (n = 16) or lamc1 morphants (n = 8). h, area of cell masses in control (n = 22) or lamc1 MO-injected embryos (n = 2). In c, d, g and h, error bars are s.d., box the s.e.m. and grey lines are mean. n.s. is *P < 0.05

## DISCUSSION

Altogether, our results suggest that the mechanical deformation that invading transformed cells experience while migrating through confined environments promotes their trans-differentiation into mesenchymal cell types. DNA damage causes a partial EMT phenotype in breast cancer cells by the upregulation of snail1 [18]. Our data implies that KRas^V12^-invading cells experience DNA damage as they migrate, and our previous work showed that a fraction of transformed cells express *snail1b* in our system [7]. Altogether, these results suggest that DNA compression and possibly damage as invading cells migrate could promote trans-differentiation of KRas^V12^ cells into mesenchymal phenotypes. Interestingly, increasing or decreasing the confinement that transformed cells experience only affects their trans-differentiation into mesenchymal but not neural-like cells. It is not clear what impacts the fate of invaded transformed cells with neuronal morphologies. Our previous studies suggested that they can sometimes migrate along normal neurons so local environmental signals may instead contribute to the trans-differentiation of these rarer cell types. Additionally, other internalized KRas^V12^/ p53^-^ cells may differentiate into different unidentified cell types, making it unclear and formation of distant metastases in two steps: first, apically-localized epithelial determinants are pinched off during BCE to de-differentiate epithelial cells (Fig. S2a), second, confinement pressure from cells migrating through tight spaces causes the de-differentiated cells to differentiate into mesenchymal cell that contribute to distant metastases (Fig. S2b).

**Fig. S2:**
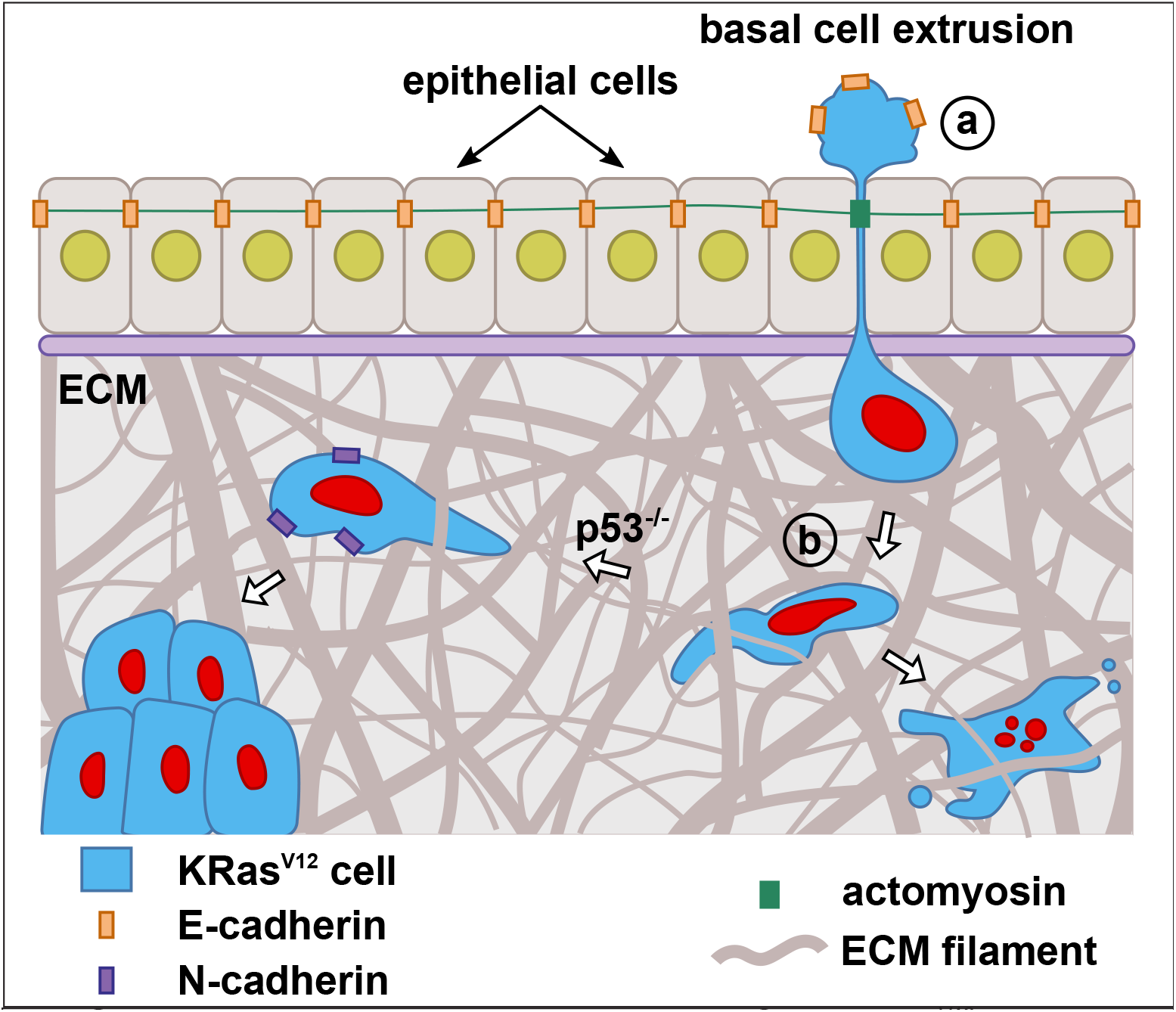
A model for cancer cell invasion by BCE. **a**, KRas^V12^ cells invade from the epithelium by BCE, losing E-cadherin and other epithelial determinants in the process. **b**, once the cells invade, they become deformed as the migrate through confined spaces within the body. Confinement prohow confinement affects their differentiation. Based on our current and previous findings [7], we suggest that physical motes trans differentiation into mesenchymal phenotypes and the formation forces contribute to cell invasion of internal cell masses. Most cells die unless p53 is also mutated.

Furthermore, increasing or decreasing confinement within the body results in more or less internal masses, respectively, by 4.5 dpf, indicating that the mechanical forces that cells encounter as they migrate contribute to distant mass formation. Interestingly, increased confinement affects the number, but not size, of cell masses. This could suggest that compression does not affect cell proliferation but does increase oncogenicity. In our system, increasing confinement for just one day resulted in more masses over the long-term, implying that sustained confinement is not necessary for increased KRas^V12^ cell metastasis. Because most invaded cells die by 4.5 dpf in embryos with a wild-type p53 background [7], the physical forces cells encounter and nuclear damage may contribute their death. However, when p53 is absent and cell death programs are hindered, these same physical forces could instead promote their trans-differentiation into mesenchymal cell types. In this way, the confines migrating cells encounter could act as a mechanical gymnasium, selecting for survival of the most aggressive, more mesenchymal-like, cells to survive.

While our zebrafish model allows us to follow all invading cells within a single organism, it is important to acknowledge that the signals and forces may be different to those encountered in human cancers. Admittedly, zebrafish embryos may have more embryonic signalling, which could allow for more plasticity than seen in adult cancers. In zebrafish, invading transformed cells can migrate through the tight confines of somites and collagen spikes that form fins, forces they would not encounter in human organs. Yet, it is interesting that most (but not all) masses accumulate in the head, which is likely softer tissue [22].

However, despite these caveats, our findings in zebrafish may support a role for compaction forces in tumour cell trans-differentiation and survival into metastases in mammals, as multiple studies have highlighted a role for stiff micro-environments promoting cancer progression [23,24]. For example, at early stages of tumorigenesis, the pressure from tumour hyperproliferation, induces expression of the beta-catenin pathway in the surrounding mouse colon, transforming healthy cells into cancer cells [25]. On the other hand, fibrosis occurring at later stages of disease progression, increases tumour environment stiffness that also promotes EMT [26-29]. Recent discoveries have also shown that the cell nucleus can sense shape changes caused by environmental confinement, causing the cell to adapt its response to surroundings [30,31]. Our findings suggest that confinement pressure directly transforms de-differentiated epithelial cells into mesenchymal phenotypes that promote their survival into distant metastases.

## METHODS

### Zebrafish use

All animal procedures were performed according to the UK Animal (Scientific Procedures) Act 1986 and carried out under Home Office Project Licence number PPL P946C972B, which was subject to local AWERB Committee review and Home Office approval. The following zebrafish lines were used: Ekkwill, AB/Tuebingen, Tuepfel long fin, Tg(*actb1:mCherry–utrCH*) [12], Tg(*bAct:hRas-eGFP*) [32], and *lamc1*^*sa379*^ mutant (sleepy) [20]. Embryos were obtained by natural spawning and raised in E3 medium at 28.5 °C. Embryos used for imaging were transferred to E3 with 0.003% N-phenylthiourea (PTU-E3, Merck) at 24 hpf to inhibit pigmentation. Sex cannot be determined before 5 dpf in zebrafish, so was not factored here.

### Microinjections and fluorescence sorting

Transposase and H2B-RFP mRNA were *in vitro* transcribed using the SP6 mMESSAGE mMACHINE Transcription Kit (Thermo Fisher Scientific), and purified using NucAway spin columns (Thermo Fisher Scientific). 2 nL of a 10-μL injection mix, comprised of 150 ng krt4:EGFP-T2A-KRas^V12^ DNA [7], 200 ng transposase mRNA, 0.2 pmol *p53* morpholino (Gene Tools, 5′-GCGCCATTGCTTTGCAAGAATTG-3′), 1 μL phenol red (Sigma) in nuclease-free dH2O (Ambion), was microinjected into one-cell embryos. Some experiments were also injected with 50 pg H2B-RFP mRNA [33] or 0.2 mM *lamc1* morpholino (Gene Tools, 5′-TGTGCCTTTTGCTATTGCGACCTC-3′). Embryos were sorted for expression of transgenes at 1 dpf using a fluorescence dissection microscope.

### Immunohistochemistry

Immunofluorescence was performed as described previously [7]. 2 and 4.5 dpf embryos were fixed in 4% paraformaldehyde, 4% sucrose, and 0.1% Triton X-100 in PBS overnight at 4 °C. Embryos were incubated with primary antibodies overnight at 4C in 10% goat serum. Primary antibodies used were against chicken α-GFP (Abcam, 1:2000), rabbit α-phospho-Histone H2A.X (Cell Signalling, 1:100), rabbit α-p63 (GeneTex, 1:100), mouse α-E-cadherin (BD Biosciences, 1:200), mouse α-N-cadherin (BD Biosciences, 1:100 and Abcam, 1:100). Embryos were incubated with appropriate secondary antibodies (goat α-chicken-AlexaFluor-488, goat α-rabbit-AlexaFluor-568, or goat α-mouse-AlexaFluor-647 (ThermoFisher Scientific, at 1:200) in 10% goat serum overnight at 4 °C. Nuclei were stained with 20 μM DRAQ5 (ThermoFisher Scientific) for 30 min and washed twice with 0.5% PBST. Embryos were mounted in ProLong Gold (ThermoFisher Scientific) between a #1.5 glass coverslip and a microscopy slide, using electrical tape as a spacer and imaged on a Yokogawa spinning disk confocal microscope with an Andor iXon camera. Images were captured using either a dry 10x lens (NA 0.30; Nikon), a dry 20x lens (NA 0.75; Nikon) or a water-immersion 40x lens (NA 1.15; Nikon).

### Live Imaging

Live embryos were anaesthetised in MS-222 (Sigma-Aldrich), then mounted in 0.6% low-melt agarose as close as possible to the #1.5 glass coverslip within a slide chamber, covered with 0.02% tricaine in PTU-E3, and incubated in a controlled environment chamber at 28 °C. Imaging was done from 1 to 2 dpf with 7-min time intervals on a Yokogawa spinning disk confocal microscope with an Andor iXon camera with a dry 20x lens (NA 0.75; Nikon).

### Agarose confinement

To induce compression, 1 dpf embryos were dechorionated, anaesthetised in MS-222 (Sigma-Aldrich) and embedded in 2% low-melt agarose in E3. Embryos were covered with 0.02% tricaine in PTU-E3, and raised at 28.5°C until 2dpf, when agarose was removed manually by gently scraping it with scalpels. Embryos were immediately fixed using the above protocol. Some of the embryos injected for this experiment were used as controls, covered with 0.02% tricaine in PTU-E3, and raised at 28.5°C until 2 or 4.5 dpf (0% agarose embryos).

### Quantitative analysis

All our quantitative analysis was performed using SIESTA [34,35], and custom scripts written in MATLAB (Mathworks) using the DIPImage toolbox (TU Delft).

### Circularity

To quantify nuclear circularity, we used a mask manually overlaid on the nucleus edge. Nuclear circularity was determined as: where a is the nuclear area and p the perimeter. Circularity is one for circles and lower than one for non-circular shapes.

### Nuclear damage quantification

To characterize cells that presented nuclear damage in the surface of the embryo or after invading, embryos injected with EGFP-KRas^V12^/ p53MO were stained for GFP, phospho-Histone H2A.X as a DNA damage marker and E-cadherin to denote the epithelium to distinguish whether KRas^V12^ cells had invaded. The number of invaded cells that were both KRas^V12^ and phospho-Histone H2A.X -positive was quantified. Percentages were calculated as the number of KRas^V12^ cells expressing phospho-Histone H2A.X, divided by the total number of invaded KRas^V12^ cells in the surface or inside the embryo.

### Cell Density

To measure neural tube cell density, z-stacks with a z resolution of 0.5 μm of embryos expressing a membrane marker (HRas-GFP) and a nuclear stain (DRAQ5) were acquired. Then the surface of the neural tube under somites 14-15 was visually detected and a substack was created from 19 μm to 21 μm underneath. A maximum Intensity Projection (MIP) was generated and the area under the somite of interest was delineated by manually overlaying a mask. Finally, the number of nuclei were counted to quantify the cell density as:

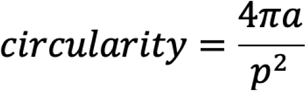

A similar procedure was used to quantify the cell density in the somites. A substack 15 to 20 μm below the surface of somites 14-15 was created and the cell density was quantified as above.

### Trans-differentiation quantification

To characterize mesenchymal trans-differentiation levels under different treatments, embryos injected with EGFP-KRas^V12^/ p53MO were stained for

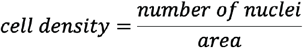

GFP, N-cadherin as a mesenchymal marker and E-cadherin or p63 to delineate the epithelium or the periderm to score only KRas^V12^ cells that had invaded beneath the periderm. The number of invaded cells that were both KRas^V12^ and N-cadherin-positive was computed. Neuron-like cells expressing EGFP-KRas^V12^ were classified based in their morphology, the expression of GFP and their location beneath the periderm. Percentages were calculated as the number of KRas^V12^ cells expressing N-cadherin or neuron morphology, divided by the total number of invaded KRas^V12^ cells.

### Fish area

To quantify the lateral area of control fish and those embedded in 2% agarose, embryos were stained with p63 to mark the periderm and obtain the embryo outline. Embryos were mounted laterally, and a z-stack of the whole embryo was acquired. MIPs of the resulting images were used to manually segment the fish outline.

### Internal cell masses

For the analysis of number and size of internal cell masses by 4.5 dpf, we imaged z-stacks of EGFP-KRas^V12^/ p53MO-injected embryos, fixed and stained with E-cadherin and GFP. To identify masses, we created binary masks from MIP of the EGFP-KRas^V12^ signal, by selecting pixels above an intensity threshold. The intensity threshold was the mean image intensity plus four standard deviations. Individual masses were labelled in the binary image and their area was quantified. Labelled objects smaller than 0.0025 mm^2^ were ignored for the analysis, to avoid including individual cells. Cells mis-expressing EGFP [7] were also excluded from the analysis. To ensure that the labelled objects were indeed internal masses we visually confirmed that the EGFP-KRas^V12^ signal was below the E-cadherin-labelled epithelia in the z-stacks.

## Statistical analysis

To evaluate sample means, we used a non-parametric Mann–Whitney test [36]. To compare more than two groups, we used a Kruskal–Wallis test to reject the null hypothesis, and a Mann–Whitney test with the Holm-Sidak adjustment for pairwise comparisons.

**Supplementary Movie 1**. An epithelial cell expressing EGFP-KRasV12 (green) invades trough BCE in an embryo expressing mCherry-Utrophin along with H2B-RFP mRNA (both in magenta). A stack was acquired every 7 min for 11 h and 47 min. Time after extrusion is shown.

## Acknowledgements

We are grateful to Jon Clarke for sharing the *lamc1* morpholino and the Tg(act-b1:mCherry–utrCH) and *lamc1*^sa379^ lines with us, to Simon Hughes for sharing the Tg(bAct:hRas-eGFP) line and to Claudia Linker for the H2B-RFP mRNA. We thank Claudia Linker and the Rosenblatt lab for useful discussions and Rachel Moore for technical help. We are grateful to Alberto Elosegui-Artola and Rachel Moore for comments on the manuscript. T.Z.-C. was supported by an EMBO Long-Term Fellowship (ALTF 1130-2018), and the European Union’s Horizon 2020 research and innovation programme under the Marie Sklodowska-Curie grant agreement No 840767. This work was supported by a Cancer Research UK Programme Grant (DRCNPG-May21\100007), an Academy of Medical Sciences Professorship APR2-1007-2, a National Institute of Health R01GM102169, and a Howard Hughes Faculty Scholar Award 55108560 to J.R.

